# Reconstitution of the DTX3L-PARP9 complex reveals determinants for high affinity heterodimer formation and enzymatic function

**DOI:** 10.1101/2021.06.14.448324

**Authors:** Yashwanth Ashok, Carlos Vela-Rodriguez, Chunsong Yang, Heli I. Alanen, Fan Liu, Bryce M. Paschal, Lari Lehtiö

## Abstract

Ubiquitination and ADP-ribosylation are post-translational modifications that play major roles in pathways like DNA damage response and infection, making them attractive targets for therapeutic intervention. DTX3L, an E3 ubiquitin ligase, forms a heterodimer with PARP9. The complex has ubiquitin ligase activity and also ADP-ribosylates the C-terminus of ubiquitin on Gly^76^. NAD^+^-dependent ADP-ribosylation of ubiquitin by DTX3L-PARP9 prevents ubiquitin from conjugating to protein substrates. By using individually produced proteins, we have studied the interaction between DTX3L and PARP9. We identify that the D3 domain (230 – 510) of DTX3L mediates interaction with PARP9 with nanomolar affinity and an apparent 1:1 stoichiometry. Our results also suggest the formation of a higher molecular weight oligomer mediated by the N-terminus of DTX3L (1-200). Furthermore, we show that ADP-ribosylation of ubiquitin at Gly^76^ is a reversible modification that can be removed by several macrodomain-type hydrolases. Our study provides a framework to understand how DTX3L-PARP9 mediates ADP-ribosylation and ubiquitination in an inter-regulatory manner.

## Introduction

Ubiquitination is a reversible post translational modification of proteins that results in the addition of a small protein called ubiquitin (Ub) to lysine residues of a target protein. Ubiquitination modifies protein properties such as stability, localization, and protein-protein interactions to name a few [1]. Conjugation of a single Ub onto substrate proteins is called monoubiquitination and this process can proceed further by the attachment of additional Ub units resulting in polyubiquitination and formation of Ub chains. Ubiquitination on substrate proteins is achieved by the consecutive action of three enzymes. Ub activating enzyme (E1) uses ATP to activate and transfer Ub to a Ub-conjugating enzyme (E2) that forms a thioester linkage with Ub (E2~Ub). Ubiquitin ligases (E3) recruit E2~Ub complex to transfer Ub onto substrate proteins.

Ubiquitination can be subject to crosstalk with other post-translational modifications such as phosphorylation [2]. There is also growing evidence of crosstalk between ubiquitination and ADP-ribosylation (reviewed in [3]). ADP-ribosylation is the attachment of ADP-ribose to proteins by enzymes that use NAD^+^ as a substrate. ADP-ribose can be transferred as a single unit [mono-ADP-ribose (MAR)] or elongated to polymers of ADP-ribose [poly-ADP-ribose (PAR)]. In humans, majority of intra-cellular ADP-ribosylation is catalyzed by PARP enzymes of the Diphtheria toxin-like ARTD family [4]. PARylation has been shown to activate E3 ligases such as RNF146 and Trip21 [5,6] and recently MARylation of Ub on Arg^42^ and Thr^66^ by bacterial enzymes of SidE family and on Gly^76^ by human DTX3L-PARP9 complex was shown to regulate ubiquitination [7–10].

In an effort to identify critical genes that determine the outcome of treatable and fatal diffuse large B-cell lymphoma, a gene named *BAL* (B aggressive lymphoma) was identified [11]. This gene was then re-named as *PARP9* as the protein C-terminus was homologous to catalytic domain of PARPs [12]. In an attempt to define molecular processes mediated by PARP9 protein, a yeast two-hybrid screen with PARP9 as a bait identified DTX3L as an interacting protein [13]. DTX3L belongs to a family of E3 Ub ligases. Deltex family comprises five members; namely DTX1-4 and DTX3L, which have similar domains in the C-terminus of the protein sharing RING and the deltex C-terminal domain [14].

In addition to lymphomas, DTX3L-PARP9 complex has been shown to be overexpressed in tumours such as prostate and breast cancer [15–17]. DTX3L-PARP9 possibly supports tumour growth by repression of IRF-1, an important transcription factor that elicits proapoptotic responses. Overexpression of DTX3L or PARP9 repressed IRF-1 promoter and knockout of either DTX3L-PARP9 lead to increased transcription from IRF-1 promoter [18].

DTX3L-PARP9 was also shown to be involved in DNA damage response as deletion of either of the genes results in increased sensitivity to DNA damage. Knockdown resulted in decrease of non-homologous end joining (NHEJ) while overexpression of DTX3L-PARP9 promoted NHEJ [19]. Furthermore, DTX3L-PARP9 was recruited to sites of DNA damage with irradiated cells [16]. DTX3L-PARP9 was also shown to be involved in anti-viral response through regulation of IFN/STAT1 pathway [20].

Biochemically, DTX3L-PARP9 was shown earlier to mono ADP-ribosylate C-terminus of Ub on Gly^76^ [19]. ADP-ribosylation at Gly^76^ is dependent on E1 and E2 processing of Ub and it was also shown that high NAD^+^ concentration promotes ADP-ribosylation of Ub and decreases Ub conjugation of substrates [19]. As an unmodified Gly^76^ is required for conjugating Ub onto substrates [21], ADP-ribosylation on this residue precludes ubiquitination. Here we describe the characterization of individually produced DTX3L and PARP9 and demonstrate that the proteins form a high affinity complex. We show that the N-terminal domain of DTX3L mediates oligomerization and that PARP9 is monomeric in solution. We also characterized the determinants of the complex formation and were able to show that the complex is mainly formed through interactions between the DTX3L D3 domain and the region of PARP9 preceding the PARP domain. Furthermore, we show that the reconstituted complex has both Ub ADP-ribosylation and ubiquitination activities. Using mass spectrometry we identified the auto-ubiquitination sites on DTX3L. We observed that the C-terminal part of DTX3L can ADP-ribosylate Ub in the absence of PARP9, but that the activity is enhanced when PARP9 is present. We also identify potential erasers for ADP-ribosylation on Gly^76^ *in vitro* using a panel of previously known ADP-ribosylation erasers.

## Materials and Methods

### Protein expression and purification

Expression vectors pNH-TrxT (#26106), pNIC-CH (#26117), pFastBac1 His-MBP (#30116) were procured from Addgene. pNIC28-MBP was constructed by replacement of TrxT with MBP [22]. Throughout the study, Uniprot IDs corresponding to DTX3L (#Q8TDB6-1) and PARP9 (#Q8IXQ6-1) were used. All constructs were verified by dideoxy sequencing. Details of expression constructs are provided in **Table S1**.

Expression constructs were transformed into BL21 (DE3) and selected on appropriate antibiotic selection plates. Overnight culture with 50 μg/ml of kanamycin were grown at 37°C. Terrific Broth-autoinduction media (Formedium, UK) supplemented with 8 g/l glycerol was used for protein expression. Briefly, when the bacterial culture grown at 37°C reached 1 O.D, the incubation temperature was reduced to 18°C overnight. Cultures were harvested by centrifugation and frozen with lysis buffer until required.

For protein expression in insect cells, recombinant bacmids were transfected into Sf21 cells using Fugene6 (Promega E2693). Supernatant containing viruses (V_0_) was harvested after seven days. These viruses were amplified once more to increase the virus titre (V_1_). Typically, 1 × 10^6^ cells were used for protein expression using an appropriate volume of V_1_ virus that was predetermined to induce growth arrest in small-scale experiments. Cell pellets were harvested 72 hours after growth arrest and frozen at −20°C with lysis buffer before use.

Cell pellets resuspended in lysis buffer (50 mM Hepes pH 7.4, 0.5 M NaCl, 10% glycerol, 0.5 mM TCEP and 10 mM Imidazole) were sonicated and centrifuged at 16,000 rpm to separate soluble material from cellular debris. Filtered supernatant was then loaded to pre-equilibrated HiTrap™ IMAC columns (Cytiva Biosciences). Columns were washed with lysis buffer, followed by wash buffer (50 mM Hepes pH 7.4, 0.5 M NaCl, 10% glycerol, 0.5 mM TCEP and 25 mM imidazole) and then eluted with elution buffer (50 mM Hepes pH 7.4, 0.35 M NaCl, 10% glycerol, 0.5 mM TCEP and 350 mM Imidazole). For constructs with MBP tag, IMAC elution was loaded into MBP-Trap columns, washed with SEC buffer (30 mM Hepes pH 7.5, 0.35 M NaCl, 10% glycerol, 0.5 mM TCEP) and eluted in same buffer with 10 mM maltose. Pooled fractions from MBP column or from other constructs without MBP tag from IMAC were digested with TEV protease (in-house preparation) [23] with 1:30 (TEV:recombinant protein) molar ratio for 1-2 days followed by reverse IMAC purification to remove TEV and cleaved tags. Proteins collected in the flowthrough were pooled and run in SEC, verified by SDS-PAGE and flash frozen and stored at – 70°C. Protein identity was verified by MALDI-TOF analysis. Co-expressed DTX3L-PARP9 complex was produced as previously reported [19].

### aSEC studies

Superdex 200 10/30 increase equilibrated with running buffer (30 mM Hepes pH 7.5, 150 mM NaCl, 5% glycerol, 0.5 mM TCEP) at a flowrate of 0.3 ml/min was used. BioRad gel filtration calibration kit was used as a standard. Fluorescent labelling was done using DyLight 488 (Thermo) at 9.8 mM. The dye was added to PARP9 FL and PARP9 M1 in an equimolar ratio and incubated for an hour at room temperature kept from the light. Excess dye was removed with a Zeba™ 7 K MWCO spin desalting column (ThermoFisher) equilibrated with running buffer.

### Ubiquitination assays

Ubiquitination reactions for DTX3L constructs were prepared in 50 μL volume by mixing (Ube1, E1) 0.4 μM, (Ubc5Ha, E2) 2 μM, DTX3L (0.67 μM) and Ub (45 μM). Tris buffer (1 M, pH 7.5) containing ATP (40 mM), MgCl_2_ (100 mM) and DTT (40 mM) was added in a 1:20 buffer to reaction volume ratio to initiate ubiquitination. Upon ATP addition, the reaction was incubated at room temperature for up to 2 h. The reaction was quenched by adding 2x Laemmli buffer at times 0, 15, 30, 60 and 120 min and analyzed using SDS-PAGE and coomassie staining. WT ubiquitin was produced in-house while single lysine ubiquitin and K0 ubiquitin were obtained from LifeSensors (Cat. # SI210).

### Identification of Ubiquitination sites

Ubiquitination reactions with DTX3L FL were conducted and resolved with SDS-PAGE. Bands corresponding to modified and unmodified proteins were cut from the gel and washed with 190 mL 40% acetonitrile (ACN) 50 mM ABic buffer until the gel piece was colourless. Then the sample was incubated at room temperature for 60 min with 100 μL of 40% ACN 50 mM ABic buffer and 2 μL 1 M DTT. After adding 4.2 μL of iodoacetamide (IAA), the sample was incubated for 30 more minutes. Following the incubation, gel pieces were washed with 190 μL 40% acetonitrile (ACN) 50 mM ABic buffer and immediately after with 190 μL of sterile water. Liquid was removed from the gel and afterwards 5 μL of trypsin (1/50 T7575, Sigma Aldrich) were added incubating the gel overnight at room temperature.

Trypsin-digested proteins were analyzed by LCMS analysis using an Easy-nLC 1000 system (ThermoFisher Scientific) coupled to a Fusion Lumos Tribrid mass spectrometer (ThermoFisher Scientific). Peptides were trapped on a Symmetry C18 0.18 x 20 mM trap column (Waters) and separated on a Waters M/Z Peptide BEH C18 130 Å 1.7 μm 0.075×150 mm column using a gradient from 97% A (0.1% formic acid) to 35% B (0.1% formic acid in ACN) over 60 min at a flow rate of 0.3 μl/min. The instrument was operated in 3 sec cycles where the MS spectra were recorded with the orbitrap analyzer at resolution of 120000 allowing the collection of up to 4e^5^ ions for maximal 100 msec before switching to MS/MS mode. Multicharged ions (threshold 2.5e^4^) were fragmented by HCD (30% collision energy) using quadrupole isolation with 1.6 Da width and 15 sec dynamic exclusion. Up to 5e^4^ fragment ions were collected for max 100 msec and analyzed in the orbitrap analyzer at a resolution of 15000. Ions between 1000 and 25000 in intensity were fragmented in the same way but MS/MS scans were recorded in the ion trap (rapid mode) aiming at higher sensitivity (threshold 1e^4^, max 300 msec).

Raw data were and analyzed with Proteome Discoverer 2.2 (ThermoFisher Scientific) using SEQUEST as search engine. MS/MS spectra recorded in the ion trap were processed with a 0.6 Da mass tolerance and data recorded with the orbitrap with a 0.02 Da tolerance. Raw data were recalibrated with the SWISS-PROT human database (9605, v2017-07-05) and after spectrum selection searched against the same database with the following settings: precursor mass tolerance 10 ppm, trypsin cleavage with up to 2 missed cleavages, carbamidomethyl on Cys, oxidation on Met, deamidation on Gln and Asn, protein N-terminus acetylated, and GG adduct on Lys as optional modifications.

Ubiquitin remnant peptides (GG peptides attached to lysine) were identified by searching for +114.042928 Da in lysine residues. Only peptides that showed 100% abundance in ubiquitinated sample when compared to the control sample were taken as ubiquitinated peptides. As trypsin cannot cleave at Ubiquitin modified lysines, peptides with C-terminal K-GG were excluded.

### ADP-ribosylation assays

The assays were performed in 20 μl of volume containing 50 mM Tris-HCl (pH 7.5), 50 mM NaCl, 5 mM MgCl_2_, 1 mM DTT, 0.5 mM ATP, 0.1 mg/ml His6-T7-ubiquitin, 12.5 mM biotin-NAD^+^,(−/+ unlabeled β-NAD^+^), 10 mg/ml each of Ube1 (E1) and UbcH8 (E2), and 0.25 mg DTX3L alone, 0.05 mg His6-DTX3L(557-740), 0.5 mg PARP9 alone, 0.25 mg DTX3L + 0.5 mg PARP9 mixture (pre-incubated at 4°C for >60 min to form complex), 0.2 mg recombinant DTX3L-(His6)PARP9 purified from bacteria, with or without 2 mg Histone H2A. The reactions were incubated at 30°C for 30 min, quenched by adding 2x Laemmli buffer, separated in SDS-PAGE, and followed with detection by Neutravidin-DyLight-800 (Pierce 22853) for biotin-ADP ribosylated Ub and anti-Ub-H2A (Cell signaling Technology #8240) for H2A ubiquitylation.

### Hydrolysis assays

His6-T7-Ub was ADP-ribosylated in the presence of 1 mM unlabeled β-NAD^+^. The unreacted β-NAD^+^ was removed with Zeba Spin Desalting column (Thermo Scientific #89892). The hydrolyses of ADP-ribose (ADPr)-Ub were performed in 20 mM Tris-HCl (pH 7.5), 50 mM NaCl, 5 mM MgCl_2_, 2 mM DTT, 0.2 mg/ml BSA and 0.3 mM hydrolase at 37°C for 60 min. ARP (Biotin-containing reagent, Cayman Chemical # 10009350) modification of the remaining ADPr-Ub were carried out in the presence of 1 mM ARP in 0.5 mM acetic acid at room temperature for 60 min [19]. The reactions were stopped with 2x Laemmli buffer mixed with neutralizing NaOH, separated by SDS-PAGE, and quantified by Neutravidin-DyLight-800 detection.

### Cross-linking and XL-MS

Cross-linking reactions were carried out at room temperature in 30 mM HEPES buffer, pH 7.5, with 350 mM NaCl, 10% (v/v) glycerol, 0.5 mM TCEP. PARP9 FL was mixed in equimolar ratios with DTX3L D3 and with DTX3L D3RD, at 15 μM and 8.6 μM respectively in 100 μL. To initiate reaction, freshly prepared BS^3^ (Thermo Scientific, #21580, Lot#OJ192037) was added to a final concentration of 500 μM to each reaction. BS^3^ was prepared by dissolving 2 mg of BS^3^ powder in 277 μL of distilled water to make a 12.5 mM working stock solution.

After incubation at appropriate times, aliquots were quenched by addition of 1 M Tris-HCl buffer, pH 8.5 (0.47 mM final concentration) to a 10 μL aliquot taken from the reaction. Samples were incubated for 20 min at room temperature and then analyzed by SDS PAGE. Bands of the expected molecular weight for both complexes (PARP9 FL-DTX3L D3 and PARP9 FL-DTX3L D3RD), were recovered from the gel for cross-linking mass spectrometry analyses.

Cross-linking of D1, D2 and D1D2 was done in a similar manner as the cross-linked complexes. Protein concentration used for these reactions was 0.5 mg/mL and quenching of the reaction was done after 10, 15, 30, 45, and 60 min. Samples were resolved using SDS-PAGE and PageBlue staining (Thermo Scientific).

Cross-linked PARP9-D3 and PARP9-D3RD were loaded on SDS-PAGE and subjected to ingel digestion. Gel bands were reduced with 5 mM DTT at 56 °C for 30 min and alkylated with 40 mM chloroacetamide at room temperature for 30 min in the dark. Protein digestion was carried out using trypsin at an enzyme-to-protein ratio of 1:20 (w/w) at 37 °C overnight.

Cross-linked DTX3L-PARP9 were digested in-solution by adding urea (6 M final concentration). The samples were reduced with 5 mM DTT at 37 °C for 60 min and alkylated with 40 mM chloroacetamide at room temperature for 30 min in the dark. Protein digestion was carried out using Lys C at an enzyme-to-protein ratio of 1:75 (w/w) at 37 °C for 4h. After diluting to 2 M urea, the digestion was continued with trypsin at an enzyme-to-protein ratio of 1:100 (w/w) at 37 °C overnight. The samples were cleaned up with C18 stageTip.

LC/MS analysis was performed using an UltiMate 3000 RSLC nano LC system coupled online to an Orbitrap Fusion mass spectrometer (Thermo Fisher Scientific). Reversed-phase separation was performed using a 50 cm analytical column (in-house packed with Poroshell 120 EC-C18, 2.7 μm, Agilent Technologies) with a 120 min gradient. Data analysis was performed using pLink 2.3.9 with the following parameters: tryptic in silico digestion with minimum peptide length=6; maximal peptide length=60 and missed cleavages=3; fix modification: Cys carbamidomethyl=57.021 Da; variable modification: Met oxidation=15.995 Da; BS^3^ cross-linker=138,068 Da. Crosslinks that appeared at least in two of the four measurements were taken for further analysis. XL-MS figures were generated using xiNET [24].

### Biolayer interferometry (BLI)

Determination of the affinity parameters was performed using the Octet RED384 (Forté Bio). All binding studies were performed in freshly prepared 30 mM HEPES buffer, pH 7.5, containing 150 mM NaCl, 5% (v/v) glycerol, 0.5 mM TCEP, 10 mg/ml BSA and 0.05% (v/v) Tween 20. The buffer was filtered with a 0.22 μm Minisart® PES syringe filter (Sartorius).

PARP9 FL and heat-inactivated BSA were biotinylated with 1 mM NHS PEG biotin. Biotinylation reaction was incubated for 30 min at room temperature and the excess biotin was removed with a Zeba™ 7k MWCO desalting column (Thermo Scientific).

The affinity measurements were carried out at 30°C using a working volume of 200 μL in black Greiner 96-well flat-bottom plates and 1000 rpm agitation. Eight streptavidin SA Dip and read ™ biosensors (FortéBio) were loaded with biotinylated 20 μg/mL BSA as a reference for nonspecific interactions. Eight streptavidin SA sensors were loaded with 10 μg/mL biotinylated PARP9 FL. Binding affinity was tested with DTX3L D3RD and DTX3L FL at 320, 160, 80, 20, 5, 2.5, 0.625 and 0 nM diluted in the working buffer. The association step was kept for 30 min to ensure signal saturation.

FortéBio Data Analysis HT 11.0 software was used for data analysis. After subtracting the signal from the reference sensors from the signal of the sensors loaded with PARP9 FL, the binding sensogram was aligned at the beginning of the association step. The sensograms were fit using a global model assuming a 1:1 interaction model in steady state.

### SAXS data collection and analysis

DTX3L FL (3.9 mg/mL) and D1D2 (17 mg/mL) were analyzed by SEC SAXS at Diamond Light Source B21 beamline (Oxfordshire, United Kingdom). Samples were run by injecting 60 μL to a Superose 6 increase 3.2/300 column at a flowrate of 0.6 mL/min with 30 mM Hepes (pH 7.5), 350 mM NaCl, 5% glycerol, 0.5 mM TCEP. The column was connected to the SAXS system on the B21 beamline. Based on the Guinier Rg, matching frames from the main peak were averaged. Data processing was done using ScÅtter [25] to determine the maximum distance (Dmax), volume and Rg. Molecular weight was determined with Primus (Atsas suite 3.0.3) using the MW and I(0) of BSA [26]. Additionally, MW was also calculated with the intensity file with the SAXSMoW server [27]. Data for DTX3L FL could not be processed due to the lack of uniformity in the data recorded.

## Results

### PARP9 is a monomer in solution

Computational predictions on human PARP9 indicate three structural domains in PARP9 (**Fig. 1A**). In order to characterize full-length PARP9, we expressed and purified PARP9 from insect cells. Several truncations were also produced to study individual domains (**Fig. 1A** and **Table S1**). Analytical size exclusion chromatography (aSEC) studies show that full-length PARP9 elutes at a volume corresponding to that of a monomer (**Fig. 1B**). Further studies with multi angle light scattering (MALS) also confirmed that PARP9 exists as a monomer with an observed molecular weight of 94 kDa with a theoretical molecular weight of 96 kDa (**Fig. 1C**).

**Fig 1.**
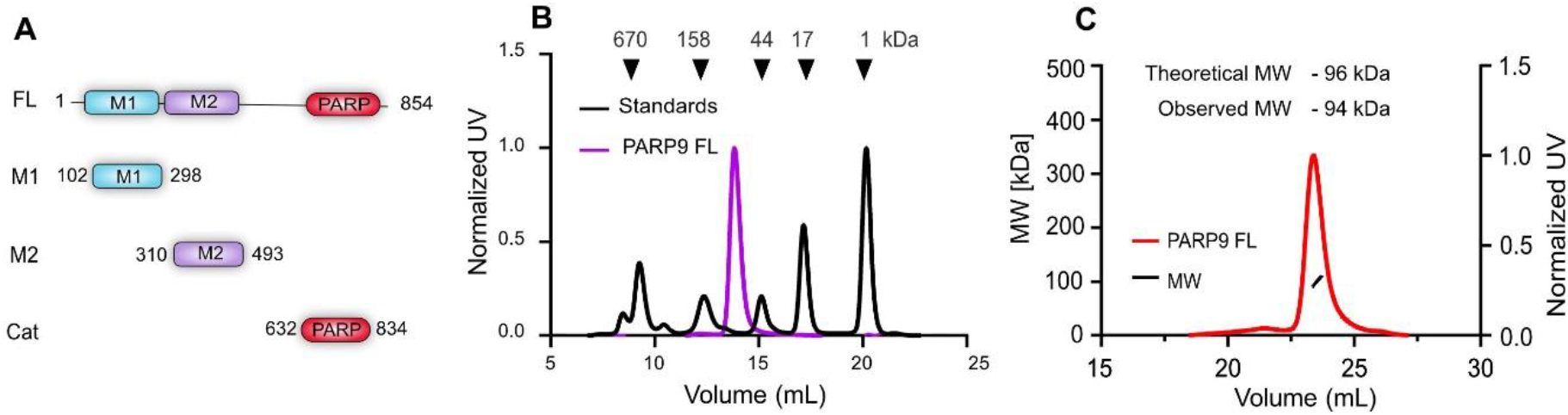
PARP9 domains and solution behaviour. A) Domain organization of PARP9 and constructs used in the studies. B) aSEC analysis of FL PARP9. Protein standards eluting at expected volume are indicated by black arrows and their corresponding molecular weight. C) SEC-MALS analysis of full-length of purified PARP9.

### D2 domain of DTX3L is the major determinant of oligomerization

Full-length DTX3L and several truncations were produced from insect cells and *E. coli* respectively (**Fig 2A** and **Table S1**). Interestingly, full-length DTX3L appeared to be oligomeric based on the protein eluting at a volume corresponding to molecular weight higher than 158 kDa with a theoretical mass of 84 kDa in aSEC. (**Fig. 2B**). To identify the domain responsible for DTX3L oligomerization, several deletion constructs were produced. We observed that the N-terminal domains spanning regions D1-D2 were oligomeric with an elution volume close to 158 kDa, contrary to an expected value of 23 kDa (**Fig. 2B**). In subsequent experiments with smaller D1 and D2 constructs, D2 domain appeared oligomeric with an elution volume that corresponds to proteins higher than 44 kDa as opposed to a theoretical 11 kDa (**Fig. 2B**). The behaviour of D1 domain seemed monomeric as its elution profile coincided with a theoretical value of 11 kDa (**Fig. 2B**). Cross-linking experiments with BS^3^ showed that D1-D2 forms an oligomer of about 125 kDa, a value that lies between the expect MW of a pentamer (113 kDa) and hexamer (136 kDa) (**Fig. 2C**). D1 domain alone showed a small amount of dimer formation and D2 domain showed bands corresponding to a tetramer (**Fig. 2D,E**). To further characterize DTX3L oligomer formation, we used small angle X-ray scattering (SAXS) with purified D1-D2 domains and full length DTX3L (data not shown for latter). We observed that the elution peak for D1-D2 domains was not entirely symmetric suggesting the presence of multiple species (**Fig. S1A**). On the other hand, intensity measurements for FL DTX3L indicate a polydisperse sample, which prevented the analysis of the sample. The relatively low uniformity observed agrees with the cross-linking data and the calculated MWs based on two different methods (114 kDa from PRIMUS and 121 kDa from SAXSMoW) fit well with the calculated MW of a pentamer but does not exclude possibility of a hexamer (**Fig. S1**).

**Fig 2.**
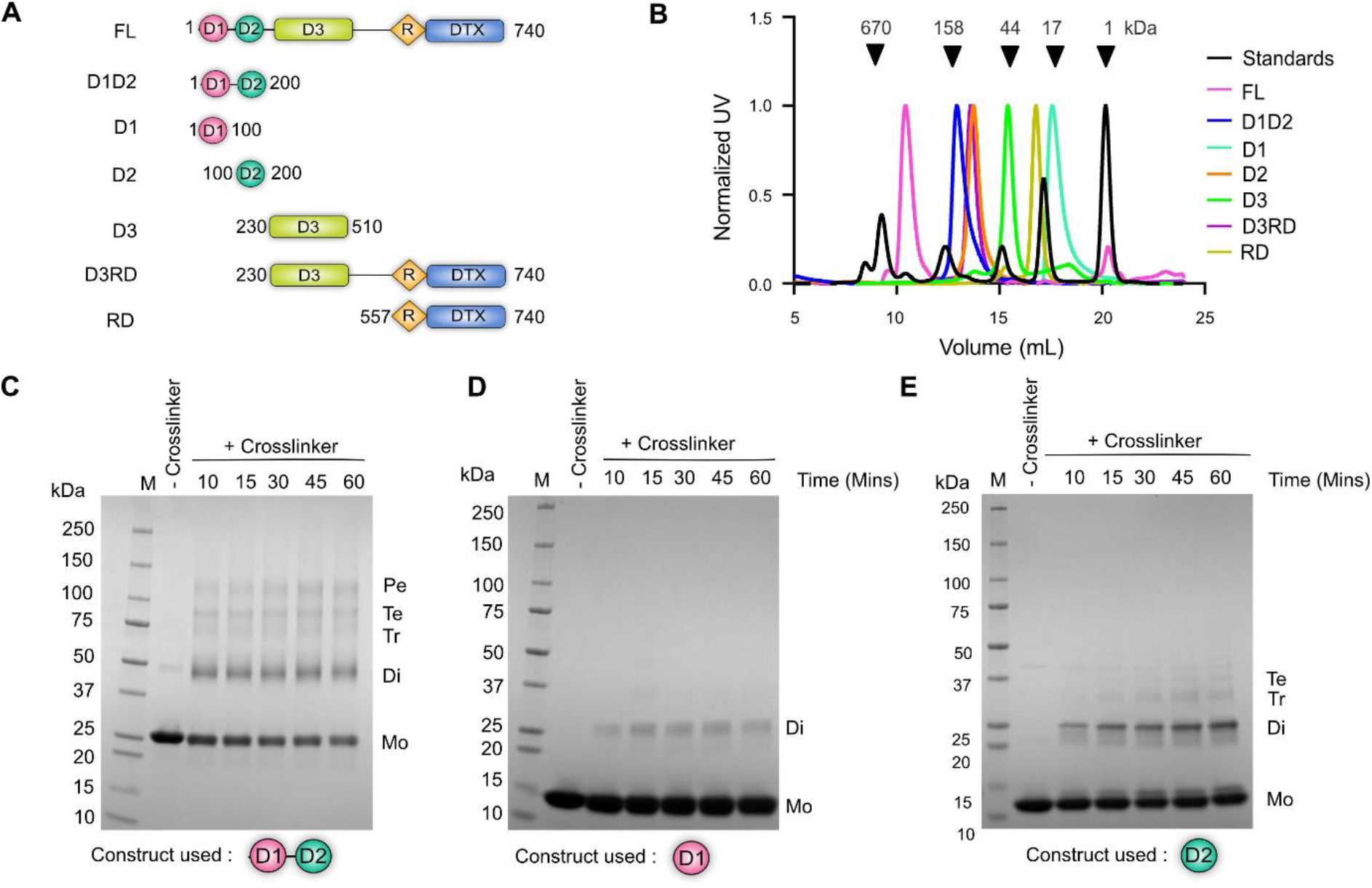
DTX3L D2 domain is the major mediator of oligomerization A) Domain organization of DTX3L and constructs used for experiments. B) aSEC of purified DTX3L proteins. Elution volumes of gel filtration standards are indicated by black arrows and their corresponding molecular weights. C, D, E) SDS-PAGE showing crosslinking analysis of purified proteins with BS^3^ for D1-D2, D1 and D2 constructs, respectively.(Abbreviations on the sides of the gel (Mo monomer, Di- dimer, Tr- trimer, Te- tetramer, Pe- pentamer).

### D3 domain is sufficient for interaction with PARP9

With purified proteins in hand, we then analyzed complex formation between full-length PARP9 and various DTX3L constructs. We used size exclusion chromatography coupled with fluorescently-labeled PARP9, which enabled us to distinguish DTX3L-PARP9 complexes from DTX3L oligomers. As FL DTX3L elutes at an early volume corresponding to a high molecular weight protein, complex formation with FL PARP9 does not significantly change the elution volume. Therefore, we labelled PARP9 with a fluorophore to aid the visualization of complex formation in size exclusion chromatography. We confirmed that chemical labelling with a fluorophore did not alter solution behavior of PARP9 (**Fig. S2**). Labeled PARP9 was then mixed with full-length DTX3L and analyzed using size exclusion chromatography (**Fig. 3A**). Fluorophore labelled PARP9 shifts elution volume when incubated with full-length DTX3L indicating complex formation. Therefore, full-length PARP9 readily formed complexes with full-length DTX3L. Subsequent experiments with DTX3L truncations revealed that D3RD (D3-R-DTX) and D3 domain formed complexes with full-length PARP9 (**Fig. 3B,C**). N-terminal D1-D2 domains and RD domains of DTX3L were dispensable for complex formation. Similarly, the catalytic domain and M1 of PARP9 were unable to form complexes in fluorescence size exclusion chromatography (**Fig. S3**).

**Fig 3.**
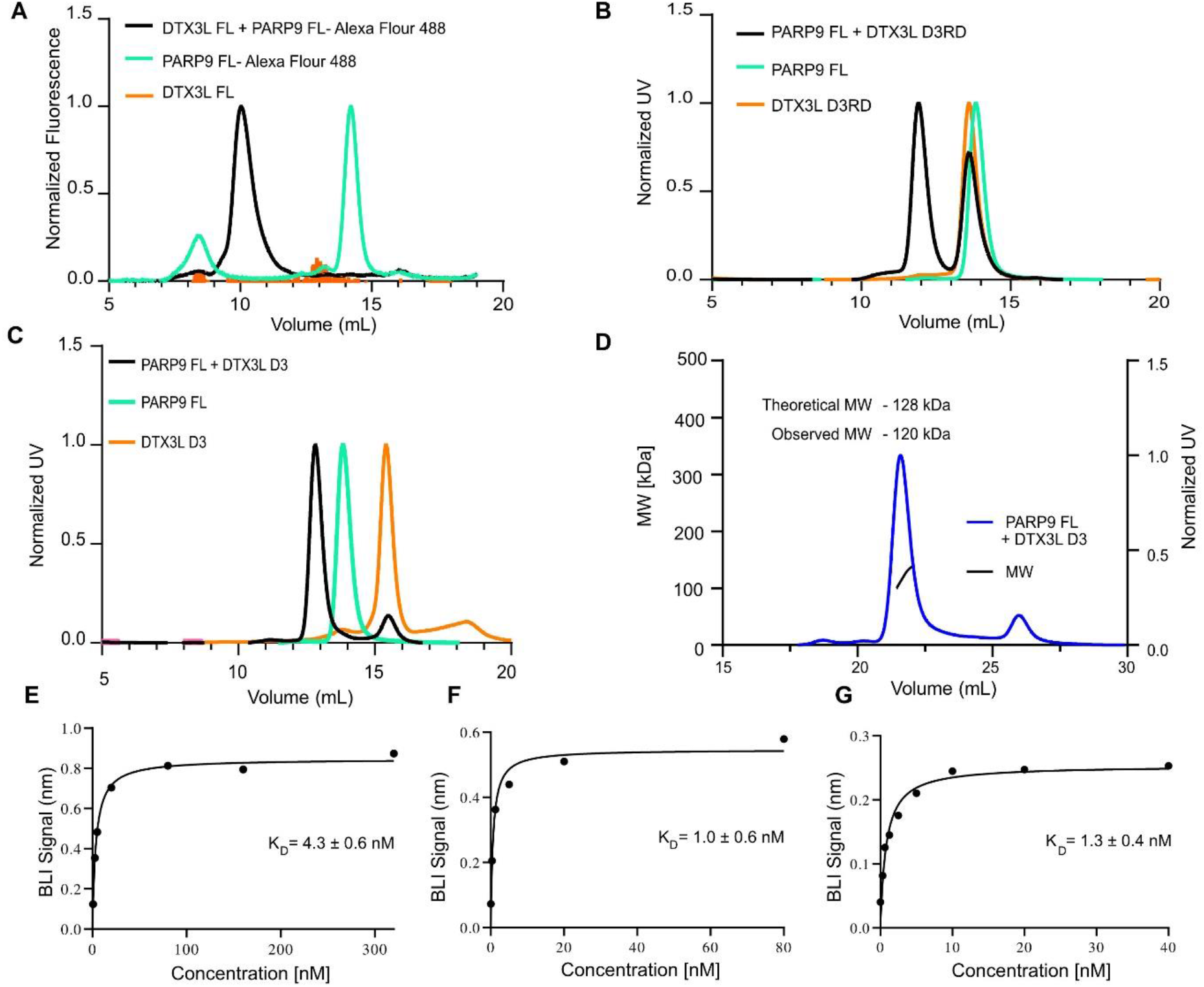
D3 domain of DTX3L is responsible for complex formation. A) Fluorescence size exclusion chromatography of full-length DTX3L and fluorophore labelled PARP9. B) SEC showing complex formation with full-length PARP9 and DTX3L D3RD domains. C) SEC showing complex formation with full-length PARP9 and DTX3L D3 domain. D) SEC-MALS analysis showing 1:1 stoichiometry of full-length PARP9 and DTX3L D3 domain. E) Bio-layer interferometry (BLI) quantitation of protein-protein interaction between PARP9 full-length and full-length DTX3L. F) Bio-layer interferometry (BLI) quantitation of protein-protein interaction between PARP9 full-length and DTX3L D3RD. G) Bio-layer interferometry (BLI) quantitation of protein-protein interaction between PARP9 full-length and DTX3L D3 domain. K_D_ values are reported as mean ± SD (n=3).

The complex of the DTX3L-D3 and PARP9 had an apparent 1:1 stoichiometry based on SEC-MALS (**Fig. 3D and Fig. S4**) and we then measured the affinity of complex formation of FL PARP9 with FL DTX3L, D3RD and D3. In all cases the biolayer interferometry showed single digit nanomolar affinities with KDs of 1.0 to 4.3 nM (**Fig. 3E-G and Fig. S5**). This is in line with the earlier observation that the complex can be extracted from the cells directly [19].

### XL-MS identifies interaction interfaces of DTX3L and PARP9

In order to get further insight into the complex formation we turned into cross-linking mass spectrometry (XL-MS), a technique where proteins or protein complexes are treated with bifunctional cross-linkers linking physically proximal regions of the protein and analyzing linked peptides after proteolysis [28,29]. We performed XL-MS with full-length PARP9 and three different constructs of DTX3L to identify both inter- and intra-molecular contacts at peptide level resolution. Despite evidence of D2 domain mediating oligomerization, we observed no cross-links between D2 domains in full-length context (**Fig. 4A**). This could be interpreted as lack of proximal lysines in the regions that oligomerize. Two cross-linked sites in DTX3L, K408 and K728 were cross-linked to themselves (i.e., a peptide containing K408 was cross-linked to K408 on another peptide), suggesting that these regions at least in context of full-length DTX3L come in proximity with each other, but this is the only direct evidence from cross-linking to an oligomer formation.

**Fig 4.**
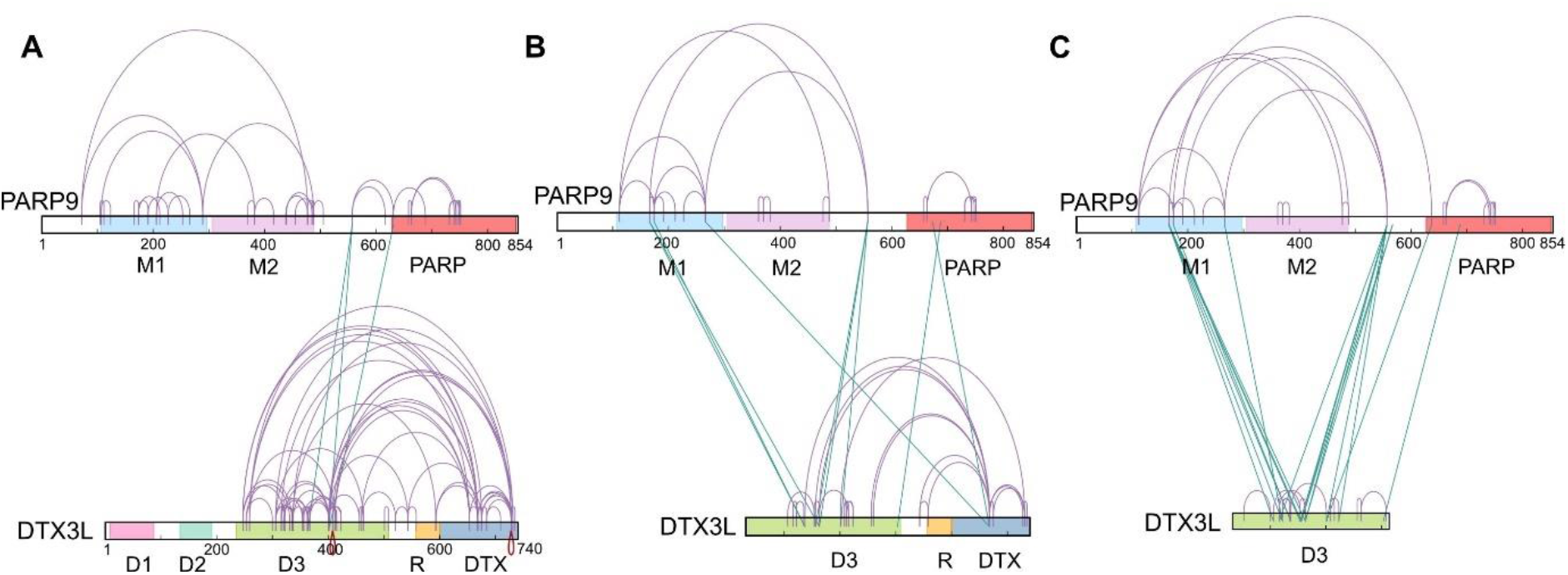
XL-MS analysis of PARP9-DTX3L complexes reveals interaction surfaces. A) Intra-protein links are shown as purple arcs. Inter-protein links are shown as cyan lines between PARP9 and DTX3L. A) Crosslink maps for full-length PARP9 and full-length DTX3L. B) Crosslink maps of full-length PARP9 and DTX3L D3RD domains. C) Crosslink maps of full-length PARP9 and DTX3L D3 domain. Intra-protein crosslinks are shown as purple lines. Inter-protein crosslinks are depicted as green lines. Symmetrical peptide cross-links (e.g., K408 cross-linked to K408 on another peptide) are shown in red lines.

For full-length proteins we observed 70 intra-protein cross-links DTX3L and 30 intra-protein cross-links in PARP9. Three inter-protein peptide cross-links between PARP9 and DTX3L were seen (**Fig. 4A**). These cross-links were observed between the D3 domain of DTX3L and the region located between M2 and PARP domains of PARP9. On the other hand, 49 high confidence intra- and inter-protein cross-links were observed between PARP9 and D3RD domains of DTX3L **(Fig. 4B)**. Of the 49 cross-links, 18 were PARP9 intra-protein cross-links, 22 DTX3L intra-protein cross-links and 9 inter-protein cross-links. With a shorter D3 domain, 21 intra-protein cross-links were seen in PARP9, 12 within D3 domain and 18 inter-protein cross-links **(Fig. 4C)**. Macrodomain M1 of PARP9 does not cross-link in the context of full-length DTX3L but steadily increases cross-linked peptides with shorter DTX3L constructs (four for D3RD and seven for D3 domain) (**Fig. 4A-C**). These results suggest that longer constructs restrain mobility of the domains with respect to each other. Two inter-protein cross-links were common for all three DTX3L constructs: PARP9 K557 is a hotspot for interaction with DTX3L D3 domain residues K401 and K363. Overall cross-linking patterns indicate that PARP9 and DTX3L form a compact structure (**Fig. 4A-C**). We used available crystal structures of PARP9 and DTX3L domains to validate the used cross-linking method (**Fig. S6**).

### DTX3L undergoes multi-domain auto-ubiqutination

We then used the purified DTX3L constructs to study auto-ubiquitination. Ubiquitination results in an increase of the molecular weight on proteins due to the covalent attachment of Ub. We show, using time course analysis, that purified DTX3L forms smears consistent with the addition of multiple Ubs **(Fig. 5A)**. All the constructs that contained the RD region (R-DTX) were active in the ubiquitination assay when E1, E2 and ubiquitin were supplied together with ATP (**Fig. 5B**). We then used mass spectrometry (MS) to map ubiquitination sites in DTX3L and were able to identify 34 auto-ubiquitination sites in multiple domains of DTX3L, with the RING domain being the only exception containing no modification sites (**Fig. 5C**). While the extent to which one or multiple ubiquitination sites in DTX3L are utilized in cells is an open question. It is probable that the protein concentration and DTX3L self-association promotes its modification *in vitro*. Ub has 7 lysine residues all of which can be ubiquitinated along with N-terminal methionine creating linear Ub chains and using MS analysis we identified that the poly-Ub chains produced by DTX3L are linked through K6, K11, K48 and K63. These linkage types indicate the potential for DTX3L to modify proteins in the contexts of proteasomal degradation and the DNA damage response (DDR) in line with previous studies [30–33] (**Fig. 5D)**. The ubiquitin reaction carried out with Ub with single lysine mutations and Ub without any lysine (K0) revealed that in all cases DTX3L gets auto-ubiquitinated consistent with multiple possible linkers and/or multiple modifications sites identified within DTX3L (**Fig. 5E**).

**Fig 5.**
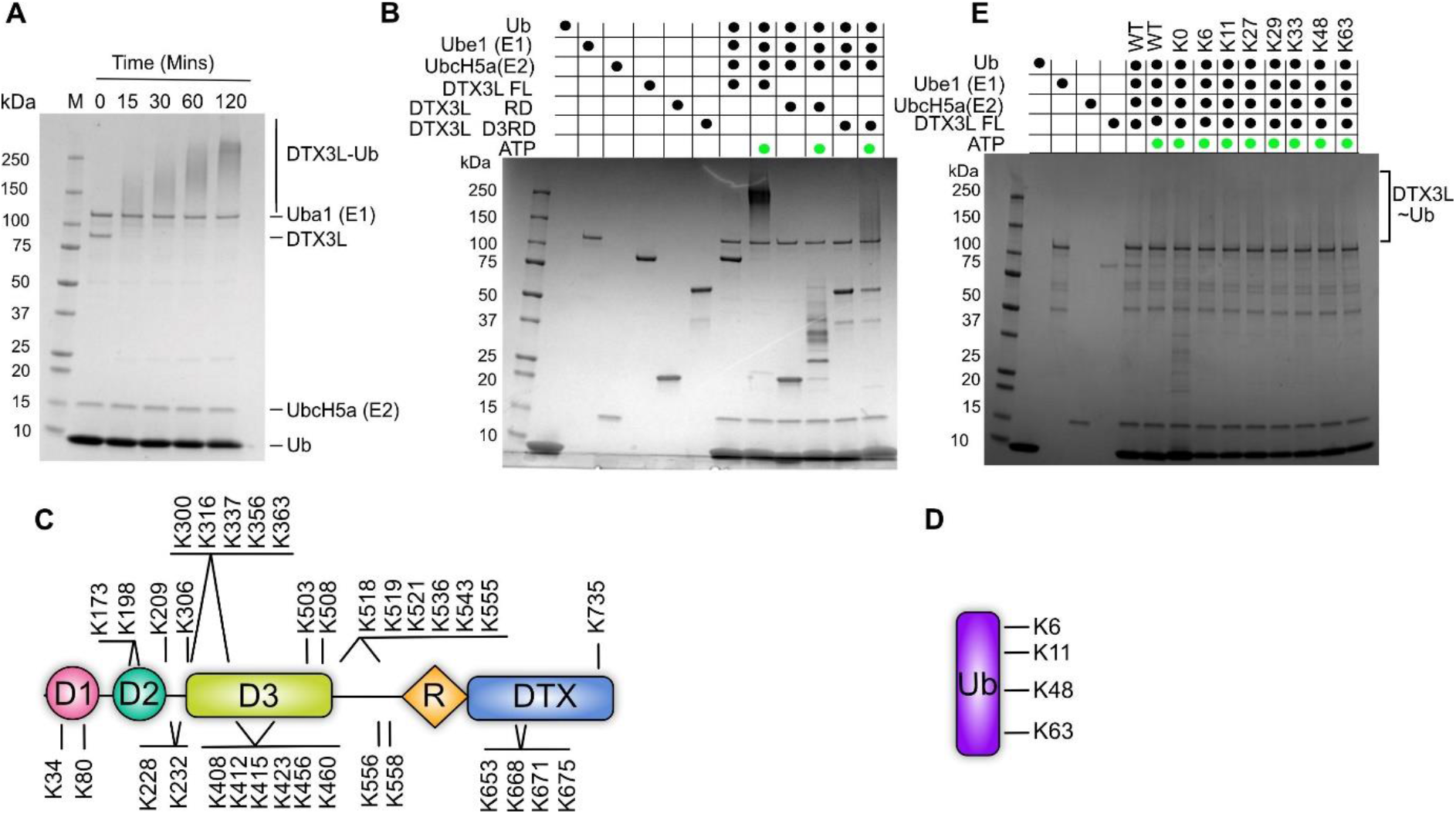
DTX3L undergoes multi-domain auto-ubiquitination. A) Time course analysis of autoubiquitnation of full-length DTX3L. Smears indicate formation of DTX3L-Ub conjugates. B) Constructs containing the RD domain are able to perform ubiquitinating activity. C) Summary of identified ubiquitination sites in DTX3L D) Mass spectrometry identified ubiquitinated lysines. E) Auto ubiquitination of DTX3L with WT Ub and lysine mutant panel of Ub.

### Heteromerization with PARP9 enhances E3 ligase and ADP-ribosyl transferase activity

In our earlier studies[19] we reported that the DTX3L-PARP9 complex mediates ADP-ribosylation of Ub on the C-terminal residue Gly^76^. Using highly pure preparations with individual proteins, we found that ADP-ribosylation of Ub Gly^76^ could be catalyzed by DTX3L alone (**Fig. 6A, lane 3**). Including PARP9 in the reaction, however, enhanced ADP-ribosylation of Ub by ~ 2.6 fold (**Fig. 6A, lane 4**). The presence of PARP9 also promoted the E3 activity of DTX3L, as indicated by the increased Ub-T7 product formation. Mono-ubiquitination of histone H2A by DTX3L increased by 60% in presence of PARP9 (**Fig 6A, lane 4**). Increasing NAD^+^ concentration to 1 mM inhibited ubiquitin conjugation as expected because ADP-ribosylation of Ub precludes conjugation to substrate as expected [19]. These data suggest that PARP9 contributes to both the ADP-ribosylation and Ub ligase functions of the heterodimer.

**Fig 6.**
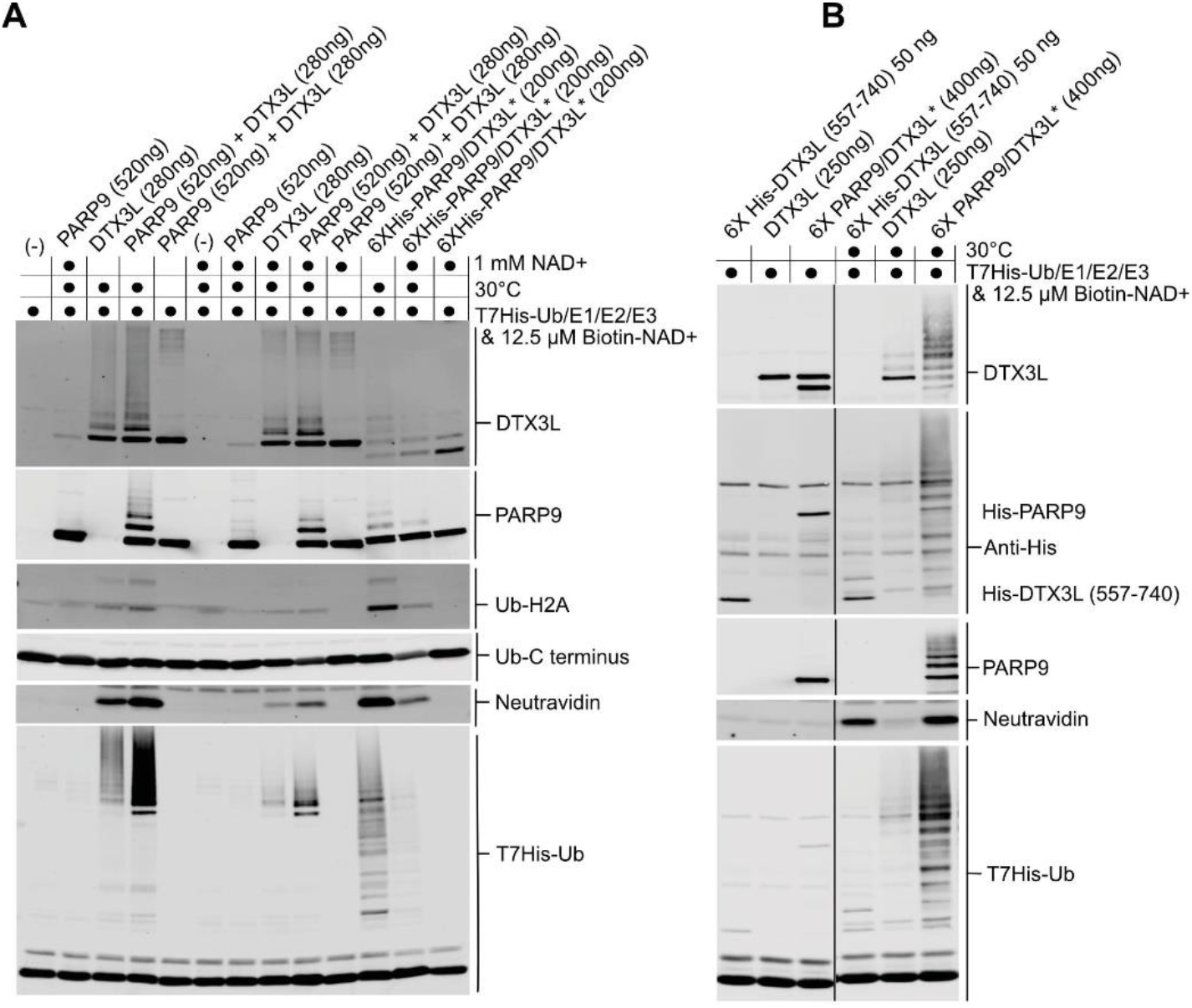
Heteromerization with PARP9 enhances DTX3L enzymatic activity. A) Assays with full-length DTX3L and PARP9, individually and after heterodimer reconstitution. Protein components concentrations are indicated. All reactions contained E1, E2, histone H2A, T7-His6-Ub, ATP, and Biotin-NAD^+^, and were incubated on ice or at 30°C. A subset of reactions was supplemented with excess NAD^+^ (1 mM). The reaction products were analyzed by immunoblotting with the indicated antibodies. ADP-ribosylation of Ub was detected using fluorescently-labeled Neutravidin, as described [19]. *Indicates co-expressed heterodimer. B) Assays comparing the DTX3L Ring-DTX domain to full-length DTX3L and DTX3L-PARP9 heterooligomer. The reactions were performed and analyzed as described in Panel A. *Indicates co-expressed heterodimer.

A ~200 amino acid C-terminal fragment of DTX3L (aa 557-740) was shown by Chatrin et al. to ADP-ribosylate Ub *in vitro* [34]. We therefore compared the activities of the DTX3L fragment to that of full-length DTX3L and the DTX3L-PARP9 heterodimer (**Fig. 6B**). The DTX3L fragment lacks the epitope for detection by the DTX3L antibody used in these assays, but was detected using 6-His antibody. We found that the fragment displays Ub (ADP-ribosyl)transferase activity that is higher than that of the full-length DTX3L. Full-length DTX3L alone is inefficient in Ub ADP-ribosylation, but the activity of the DTX3L-PARP9 heterodimer is at the same level as for the isolated DTX3L C-terminal fragment suggesting that contacts between DTX3L and PARP9 enable the ADP-ribosylation activity of DTX3L.

### ADP-ribosylhydrolases catalyze the removal of ADP-ribose from Gly^76^ Ub

During the completion of this study, it was reported that Ub Gly^76^ ADP-ribosylation can be reversed *in vitro* by deubiquitinases (DUBs) [34]. DUBs are not, however, the typical erasers of ADP-ribosylation and three families of ADP-ribosylation erasers have been reported, which is why we studied the possibility that hydrolases remove the modification. The erasers of ADP-ribosylation are grouped in the ARH family of enzymes (comprising ARH1-3), macrodomain containing enzymes (PARG, TARG1, MacroD1 and MacroD2), and nudix hydrolases. Apart from PARG, all erasers are dependent on the amino acid linkage to ADP-ribose. For example, ARH1 targets N-glycosidic bonds on Arginine, whereas ARH3 targets *O*-glycosidic bonds on serine [35–37]. MacroD1, MacroD2 and TARG1 target ADP-ribose conjugated to acidic residues like glutamic acid and aspartic acid [38–41]. In contrast to other erasers, nudix are considered as partial hydrolases since they cleave the phosphodiester bond between adenosine and ribose instead of removing complete ADP-ribose units [42].

We analyzed a panel of recombinant ADP-ribosylation erasers for the ability to remove the modification. Our results indicate that most of the enzymes tested can remove ADP-ribose from Ub- Gly^76^ (**Fig. 7A,B**). Both ARH1 and 3 had similar activities, but substrates for ARH2 remain unknown and it did not show any activity towards Ub Gly^76^ ADPr. On the other hand, amongst enzymes that contain a macrodomain fold, PARG, MacroD1 and D2 showed high hydrolysis activities.

**Fig 7.**
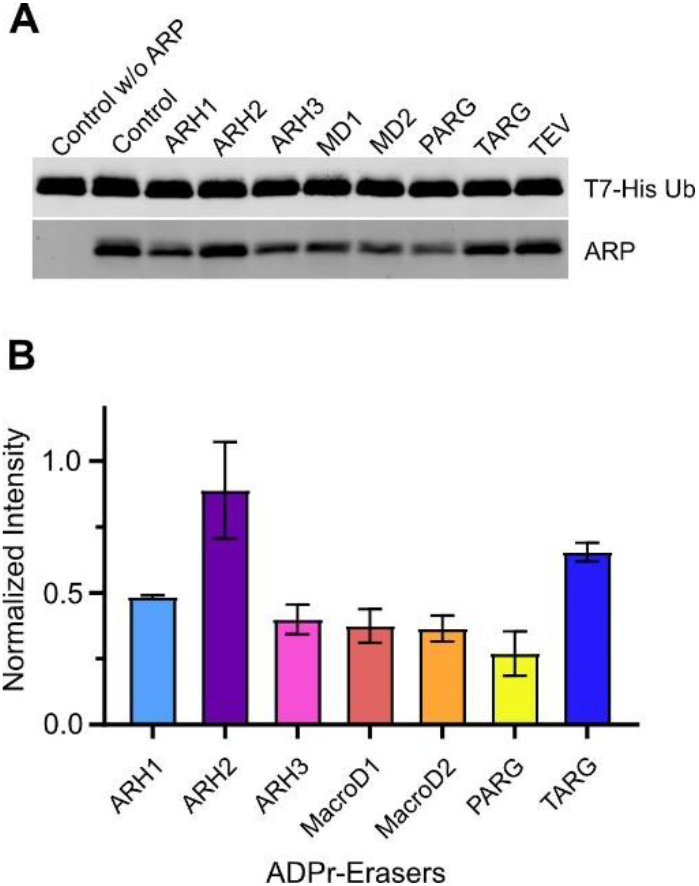
Several ADPr hydrolases can remove ADP-ribose from Gly^76^ Ub. A) Hydrolysis of ADP-ribose from the C-terminus of Ubiquitin by a panel of human (ADP-ribosyl)hydrolases. TEV protease was used as a control. (n = 2) D) Quantification of the hydrolysis activity from gel shown in A. Control Lane with ARP detection was normalized to 1.0.

## Discussion

DTX3L-PARP9 complex has both Ub E3 ligase and ADP-ribosylation activities. The significance of studies on this complex relies on its role in DNA damage repair pathways, cancer biology and in viral infection. For the first time, we have successfully purified full-length DTX3L and PARP9 and several truncation constructs. While DTX3L-PARP9 was shown to exist as a large complex in earlier studies, these studies used crude lysate to probe oligomeric status of the complex. In crude lysate preparations, other interacting proteins could modify solution behaviour of proteins. We show conclusively using pure preparations of PARP9 that it exists as a monomer in solution and DTX3L exists as a multimer. Studies with purified D1-D2 domains showed that they mediate oligomerization in line with previous studies [20]. Further dissection of these domains separately into D1 and D2 indicated that majority of the oligomerization is driven by D2 domain with minor contributions from D1 domain. It must be noted that D1-D2 domains present a heterogeneous oligomer behaviour where the dominating species in solution seem to be either a pentamer or a hexamer as calculated from SAXS data. Multimerization is not an uncommon feature of E3 ligases, with well-characterised examples of homo-dimerization (such as RNF4, BIRC7, IDOL) as well as hetero-dimerization (including BRCA1-BARD1 and MDMX-MDM2) [33,43–46]. However, dimerization of those ligases is mainly mediated by the RING domain and with a clear effect on enzymatic activity [33,47–49]. Based on our results, RING domain of DTX3L is not involved in protein oligomerization.

We confirm earlier biochemical studies that D3 domain of DTX3L is sufficient for interaction with PARP9. D3 domain binds with nanomolar affinity to PARP9 which allows the formation of stable complexes in aSEC. Using XL-MS we have gained new information about the interaction at peptide level resolution. D3 domain makes contacts with linker region between M2 and catalytic domain of PARP9 and the catalytic domain. Intriguingly, crosslinks between two and K408 and K728 (DTX3L) residues could be identified while cross-linking full-length proteins implying that DTX domains from adjacent monomers come in close proximity in an oligomeric context.

Our results suggest that multimerization may act as a regulatory mechanism based on the MARylating activity towards Ub, which can be a result of a rigid, unfavourable conformation for the full-length DTX3L enzyme as opposed to having a monomeric C-terminal fragment of the protein. Additionally, the identification of cross-linking regions between the catalytic domain suggests a more active participation from PARP9 in the reaction (**Fig. 4**). Interestingly, other interacting regions that were experimentally identified indicate a compact structure that brings to close proximity the R-DTX region of DTX3L to the macrodomains of PARP9. Taking this into account, it is possible that the compact structure of the complex facilitates the ubiquitination of PARylated proteins in the context of DNA damage. Our assumption is supported by the linkages identified by MS on ubiquitinated DTX3L, which are those relevant to the context of DDR, supporting previous findings [16].

ADP-ribosylation of Ub on Gly^76^ was initially reported *in vitro* and later shown to be present in cells [34]. However, no functions have been ascribed to this modification. It is hypothesized that conjugation of ADP-ribose on Gly^76^ by DTX3L-PARP9 makes it unable to conjugate to proteins by the canonical pathway since the modification protects C-terminal glycine making it unavailable to ubiquitinate substrate proteins [21].

Earlier, one of our labs reported the existence of cellular factors that are capable of removing ADP-ribose from Ub Gly^76^ [19]. Based on previous studies, we assembled a panel of proteins known to remove ADP-ribose from proteins. We were able to identify some candidate proteins that can remove the ADPr moiety from ubiquitin *in vitro*. An interesting observation is that PARG is the most active of all studied proteins in removing ADPr from Ub. PARG is generally inefficient in breaking the ADPr-protein bond and much more efficient in hydrolyzing ADPr-ADPr bonds [50]. TARG on the other hand seemed inefficient by comparison despite being shown to be effective in acting upon protein-ADPr bonds [41,51]. Additionally, we observed that multiple enzymes efficiently remove ADPr from Ub-Gly^76^, including MacroD1, MacroD2, ARH1 and ARH3. All these are *in vitro* studies and the physiological relevance of these erasers would require further cell-based experiments.

In summary, we have used biochemical methods to characterize the DTX3L-PARP9 complex. DTX3L binds PARP9 with nanomolar affinity and relies on multiple inter-subunit contacts to help maintain the overall structure and biochemical activity of the complex. We provided evidence for intra-protein contacts within DTX3L, and also within PARP9. The latter included contacts both within and between the two macrodomains of PARP9, which are known to provide the complex with ADP-ribose reader function [52]. The coordinated expression and efficient assembly of DTX3L-PARP9 helps to bring together the E3, (ADP-ribosyl)transferase, and ADP-reader function into a single complex.

## Supporting information

Supplementary XLMS file

## Competing interests

The authors declare that they have no competing interests.

## Funding

The research was supported by the Academy of Finland grants 287063, 294085 and 319299 (to LL) and by an NIH grant No. R01CA214872 (to BMP), and by Leibniz Association with the Leibniz-Wettbewerbs (to FL).

## Data availability

XL-MS peptide analysis is available in the supplemental file. SAXS data is deposited to SASDB.

## Author contributions

LL and BMP conceived the research. YA, CVR and CY carried out biochemical experiments and analyzed the data. HIA carried out cloning and protein production. FL was responsible for the XL-MS measurements. YA, CVR, BMP and LL wrote the publication with contributions from all the authors.

## Acknowledgements

We thank Kalle Niemi for testing ubiquitination reactions, Ulrich Bergman for expert help with ubiquitin site mapping, Hongmin Tu for expert help with BLI, Albert Galera-Prat for expert help with SAXS data analysis, and beam line scientists at Diamond Light Source B21 beamline for mail-in data collection service. The use of the facilities of the Biocenter Oulu Structural Biology core facility, member of Biocenter Finland, Instruct-ERIC Centre Finland and FINStruct, is gratefully acknowledged. We also thank the Biocenter Oulu Sequencing Center and Proteomics and protein analysis facility. We acknowledge Drs. Opher Gileadi, Cynthia Wolberger and Scott Gradia for making plasmids available through Addgene.

## Notes

### Competing Interest Statement

The authors have declared no competing interest.

### Summary of Updates

Manuscript is expanded and improved.

