## Supplementary tables and figures for "Reconstitution of the DTX3L-PARP9 complex reveals determinants for high affinity heterodimer formation and enzymatic function"

### CONTENT

**Table S1:** Expression constructs used in recombinant protein production.

**Fig. S1:** SAXS analysis of the D1-D2 construct.

**Fig. S2:** Fluorescently labelled PARP9 elution profiles.

**Fig. S3:** Additional aSEC analysis of complex formation of DTX3L and PARP9.

**Fig. S4:** SEC-MALS experiment with DTX3L-D3 domain.

**Fig. S5:** Representative BLI sensorgrams between PAPR9 and DTX3L constructs.

**Fig. S6:** Distribution of crosslink distances in XL-MS.

**Table S1.** Expression constructs and hosts used in the production of recombinant DTX3L and PARP9.

| Construct | Amino acids | Vector | Expression system |
|---|---|---|---|
| DTX3L FL | 1-740 | pFastBac1 His-MBP (Addgene #30116) | Sf21 insect cells |
| DTX3L D1D2 | 1-200 | pNIC-MBP | *E. coli* |
| DTX3L D3 | 230-510 | pNH-TrxT | *E. coli* |
| DTX3L sD3 (insoluble) | 230-485 | pNH-TrxT | *E. coli* |
| DTX3L D3RD | 230-740 | pNIC28-BSA4 | *E. coli* |
| DTX3L RD | 557-740 | pNH-TrxT | *E. coli* |
| DTX3L D1 | 1-100 | pNIC-MBP | *E. coli* |
| DTX3L D2 | 101-200 | pNIC-MBP | *E. coli* |
| PARP9 FL | 1-854 | pFastBac1 His-MBP | Sf21 insect cells |
| PARP9 M1 | 102-298 | pNIC28-BSA4 | *E. coli* |
| PARP9 M2 | 310-493 | pNIC28-BSA4 | *E. coli* |
| PARP9 Cat | 632-834 | pNIC-CH | *E. coli* |
| PARP9 LoopCat (insoluble) | 491-854 | pNIC-MBP | *E. coli* |
| PARP9 Loop (insoluble) | 491-630 | pNIC-MBP | *E. coli* |
| Ube1 (Addgene #34965) | FL | pET21d | *E. coli* |
| Ubc (Addgene #12647) | 1-76 | pET15 | *E. coli* |

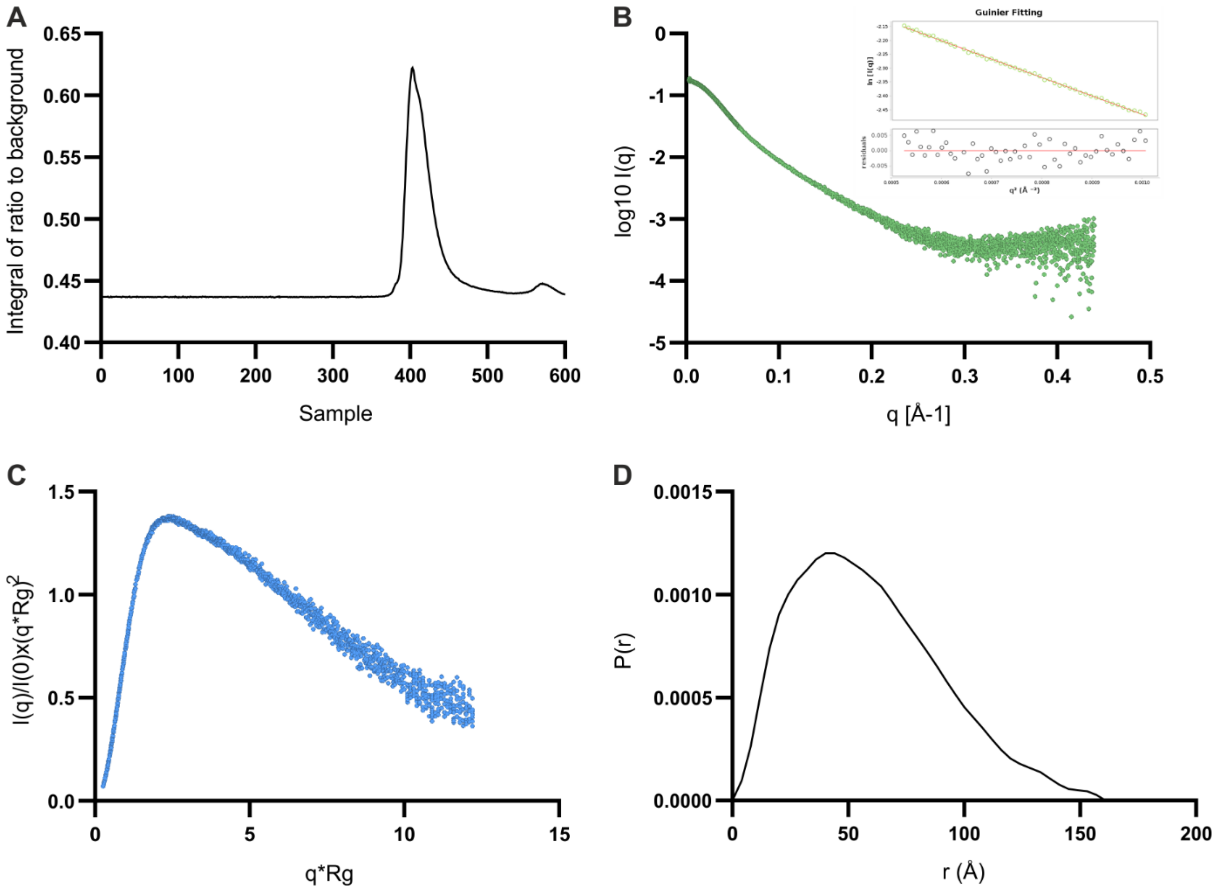


**Fig. S1.** SAXS analysis for D1-D2 domains. A) HPLC chromatogram. B) SAXS data plot with Guinier fitting plot and residuals distribution. C) Dimensionless Kratky plot analysis. D) Distance distribution analysis with a maximum distance ($D_{max}$) of 160 Å. The calculated volume is 212086 Å$^3$ for an Rg of 4.7 Å. The Porod exponent for the sample was calculated at 2.7, indicating that a mixture of flexible and compact particles.

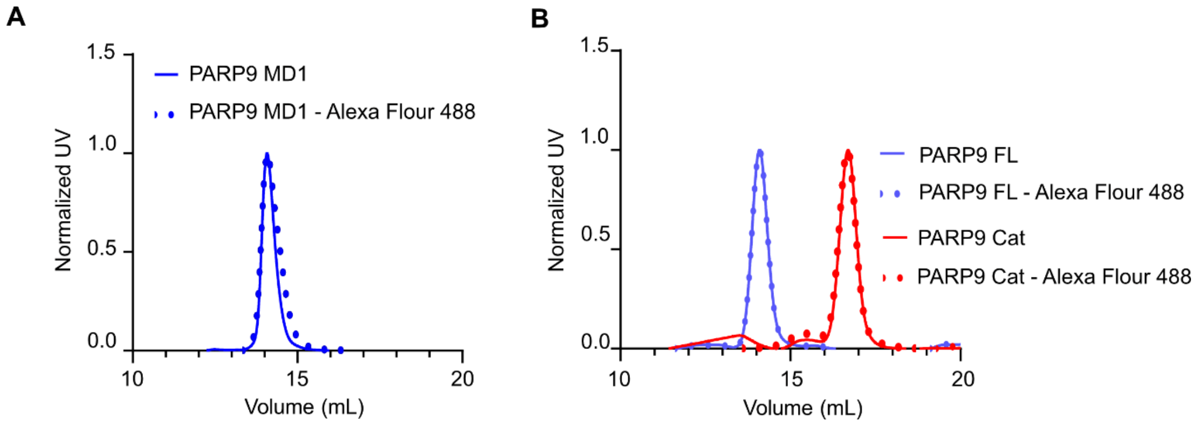


**Fig. S2.** Alexa Fluor 488 labeling of proteins does not change their SEC profile. Solid lines- unlabeled protein, dotted lines- Fluorophore labeled protein. A) PARP9 MD1. B) PARP9 FL and PARP9 Cat.

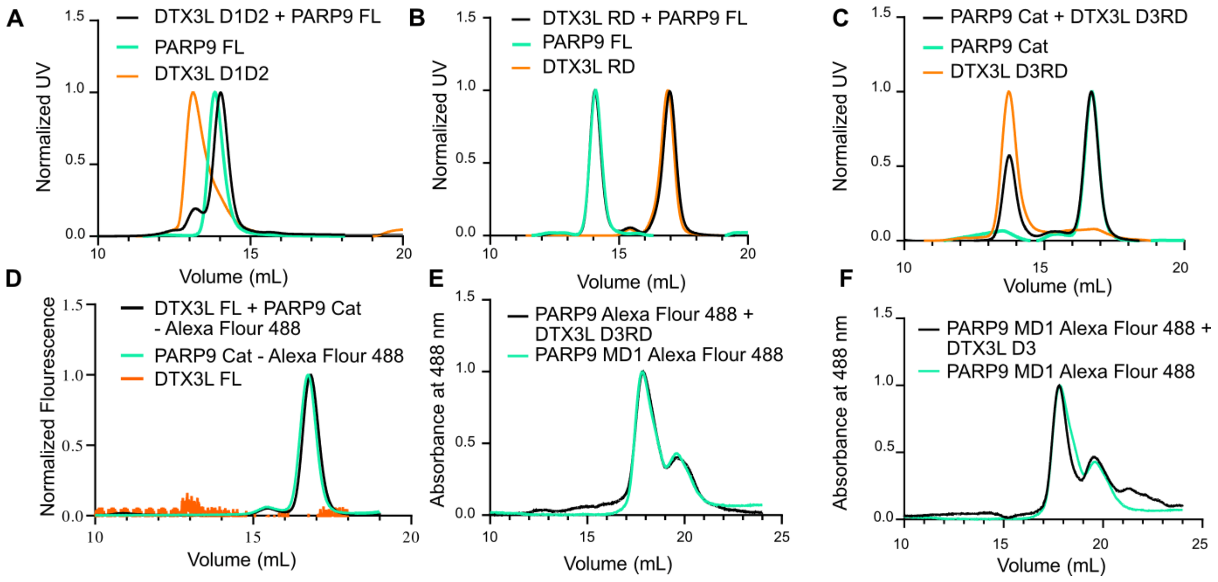


**Fig. S3.** aSEC analysis of complex formation with various constructs of DTX3L and PARP9. A) DTX3L D1D2 and PARP9 FL. B) DTX3L RD and PARP9 FL. C) DTX3L D3RD and PARP9 FL. D) DTX3L FL and PARP9 Cat. E) DTX3L D3RD and PARP9 MD1. F) DTX3L D3 and PARP9 MD1.

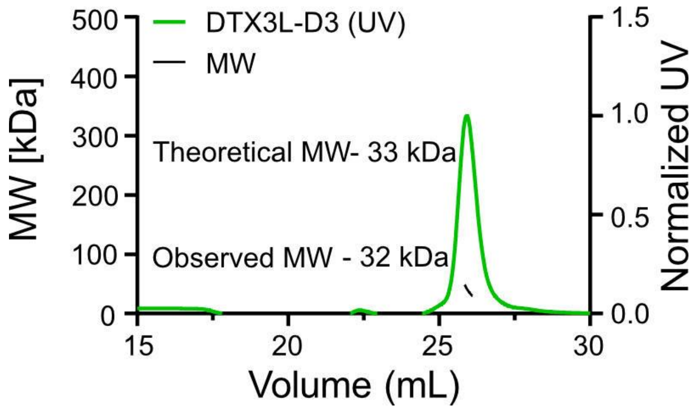


**Fig. S4.** SEC-MALS experiment with DTX3L-D3 showing that it exists as a monomer in solution.

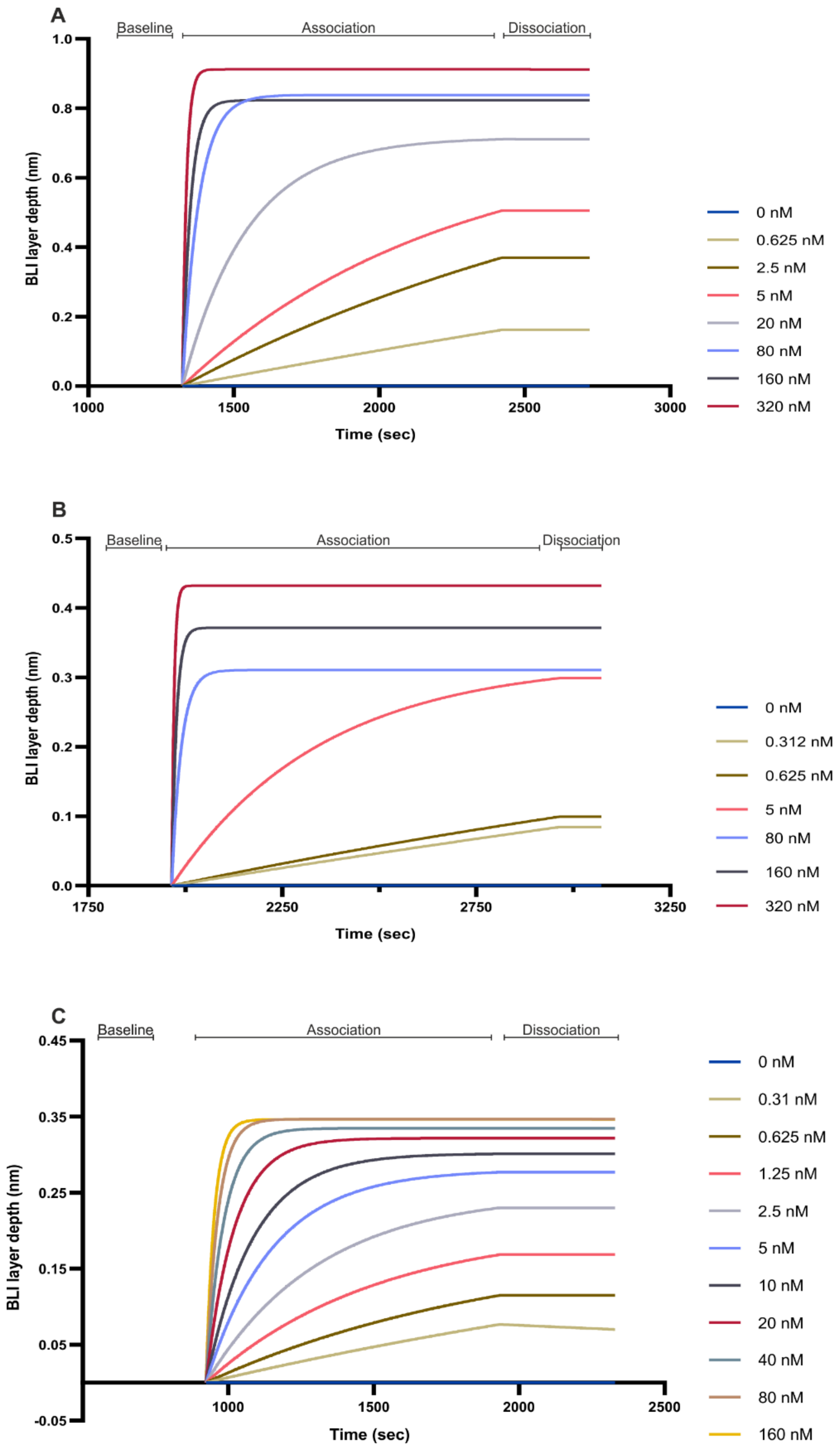


**Fig. S5:** Representative BLI sensograms between DTX3L and PARP9. A) FL PARP9 and FL DTX3L. B) FL PARP9 and D3RD. C) FL PARP9 and D3. Note that dissociation step is not taken into account for calculating equilibrium $K_D$.

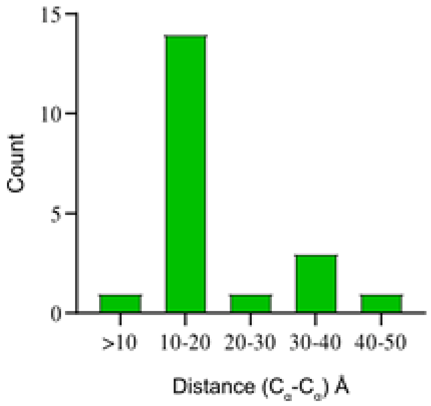


**Fig. S6.** Distance distribution of crosslinks. Number of crosslinks identified by XL-MS with the distance range (Å) within they fall. Using crystal structure of Macrodomain M2 of PARP9 (PDB: 5AIL) and modelled structures of Macrodomain M1, catalytic domain of PARP9 and Ring-DTXC structures (generated using Swiss model), we measured distances between Cα atoms of two cross-linking lysines in these structures. The majority (80%) of the cross links fall below the cutoff of 30 Å that is accepted for $BS^3$ cross-linker between two Cα atoms [1]. Two outliers in R-DTX domain could be rationalized as conformational mobility between these domains which was shown experimentally by using structural and biochemical studies [2].
